# ERAD machinery controls the conditional turnover of PIN-LIKES in plants

**DOI:** 10.1101/2025.07.05.663279

**Authors:** Seinab Noura, Jonathan Ferreira Da Silva Santos, Elena Feraru, Sebastian N.W. Hoernstein, Mugurel I. Feraru, Laura Montero-Morales, Ann-Kathrin Rößling, David Scheuring, Richard Strasser, Pitter F. Huesgen, Sascha Waidmann, Jürgen Kleine-Vehn

## Abstract

Auxin is a crucial phytohormone that regulates plant development and facilitates dynamic responses to environmental changes through subcellular control mechanisms. PIN-LIKES (PILS) are auxin transport facilitators at the endoplasmic reticulum (ER) that mediate nuclear auxin abundance and signalling. While the posttranslational regulation of PILS is important for acclimating growth responses, the molecular mechanisms involved remain largely unknown. This study demonstrates that components of the ER-associated degradation (ERAD) machinery regulate the proteasome-dependent degradation of functional PILS proteins under non-stressed conditions. We further reveal that both internal and external signals utilise the ERAD complex to differentially modulate the turnover rates of PILS proteins. Our findings uncover an additional physiological role of the ERAD complex in regulating PILS protein turnover. This finding uncovers the interplay between protein homeostasis at the ER and growth regulation, opening new molecular avenues into how plants acclimate to internal and external cues.

## Introduction

The phytohormone auxin plays a central role in coordinating plant growth responses to internal and external signals. These adaptive auxin responses largely depend on the modulation of auxin transport processes, which ensure precise distribution and availability of this hormone throughout the plant. While considerable progress has been made in understanding intercellular auxin transport, intracellular mechanisms remain less well characterised. Recent evidence suggests that subcellular control of auxin is critical for plant acclimation to environmental fluctuations (*1–6*).

The PIN-LIKES (PILS) family of auxin transport facilitators, localized at the endoplasmic reticulum (ER), plays a role in regulating nuclear auxin levels and thereby signaling (*1*, *3*, *7*). This regulation is thought to occur through a compartmentalization effect that limits auxin diffusion into the nucleus. PILS protein abundance is posttranslationally regulated by a variety of environmental cues, including light, temperature, and ER stress, as well as internal phytohormonal signals, such as brassinosteroids and auxin (*3*, *5*, *6*).

By integrating these signals, PILS proteins contribute to the acclimation of auxin-dependent growth (*2*, *3*, *5*, *6*). Despite their developmental significance, the mechanisms by which ER-resident proteins, including PILS, are regulated and subjected to degradation remain poorly understood.

The ER-associated degradation (ERAD) complex is essential for degrading misfolded or defective proteins, thereby maintaining ER homeostasis in plants (*8*). Beyond quality control, the ERAD complex is increasingly recognised to regulate the conditional degradation of properly folded proteins, often referred to as physiological clients (*9–12*). Here, we explore the role of the ERAD complex in the conditional degradation of functional PILS proteins, pinpointing a role in plant growth regulation.

## Results

Canonical PIN-FORMED (PIN) auxin carriers are predominantly located at the plasma membrane and are trafficked to the vacuole for lytic degradation (*13–15*). Given that vacuolar cargo sorting can initiate in the ER (*16*), we initially hypothesized that PILS proteins might similarly be transported to the vacuole for degradation. Functional PILS-GFP fusions (*1–3*, *5*, *6*) predominantly remain in the ER and were unaffected by Brefeldin A (BFA), a vesicle trafficking inhibitor that also disrupts vacuolar cargo delivery ((*17*), fig. S1A, B). Moreover, treatment with Vacuolar Affecting Compound 1 (VAC1), which induces the accumulation of vacuolar cargos (*18*), did not inhibit but promoted the degradation of PILS6 (fig. S1C, D). These findings suggest that, unlike canonical PIN proteins, PILS proteins do not rely on BFA- or VAC1-sensitive trafficking pathways for their lytic degradation.

To further investigate the regulatory mechanisms governing PILS protein turnover, we assessed the role of the cytosolic 26S proteasome using the pharmacological inhibitor bortezomib (BTZ) (*19*). The application of the proteasome inhibitor bortezomib (BTZ) resulted in increased fluorescence of constitutively expressed GFP-PILS3, PILS5-GFP, and PILS6-GFP in dark-grown hypocotyls compared to solvent control treatments (Fig. 1A, B). This observation suggests a general posttranslational regulation of PILS proteins. These findings strongly indicate that the turnover of PILS proteins depends on the functionality of the 26S proteasome. In addition, BTZ application notably increased the abundance of pPILS3::PILS3-GFP in apical hooks (in the *pils3-1* mutant) (Fig. 1C, D), but did not influence PILS3 expression (fig. S1E). This set of data illustrates that proteasomal degradation significantly impacts the physiological levels of PILS3 proteins. Notably, the effects of proteasome inhibition on PILS3 abundance were observed within just one hour (Fig. 1D), suggesting a rapid and likely direct mode of action.

**Figure 1:**
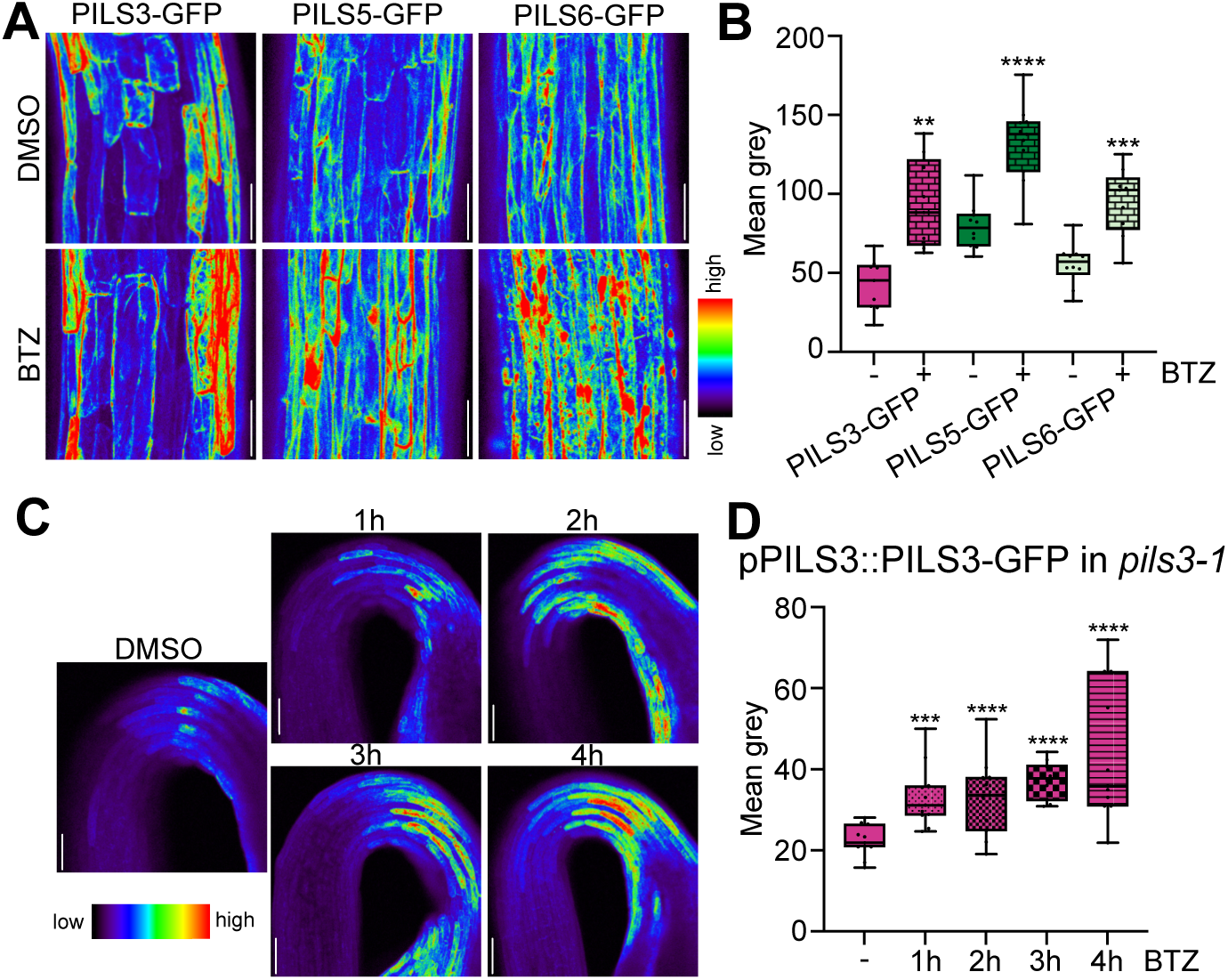
Interference with the proteasome increases PILS abundance. **A-B**, Representative images (A) and quantifications (B) of GFP-PILS3, PILS5-GFP and PILS6-GFP signal in 3-days-old dark-grown seedlings. Seedlings were grown on solid ½ MS and treated with DMSO or 40 µM BTZ in liquid ½ MS for 4h. Scale bars, 50 µm. n = 9-11, Student’s t-test between DMSO and BTZ. **C-D**, Representative images (C) and quantifications (D) of pPILS3::PILS3-GFP in the *pils3-1* background in 3-days-old dark-grown seedlings. Seedlings were grown on solid ½ MS and treated with DMSO or 40 µM BTZ in liquid ½ MS for 1-4h. n = 10, one-way ANOVA followed by Tukey’s multiple comparisons between DMSO and BTZ. In all panels with boxplots: Box limits represent 25th percentile and 75th percentile; horizontal line represents median. Whiskers display min. to max. values. P-Values: * P < 0.05, ** P < 0.01, *** P < 0.001, **** P < 0.0001. All experiments were repeated at least three times.

To molecularly elucidate the proteasome-dependent turnover of PILS proteins, we subsequently conducted an unbiased proteomic screen using antibody-based affinity purification (*20*) in conjunction with mass spectrometry. Through our proteomic analysis, we identified proteins that directly or indirectly associate with the functional GFP-tagged PILS2, PILS3, and PILS6 fusion proteins. Notably, components of the endoplasmic reticulum-associated degradation (ERAD) complex, including HMG-CoA Reductase Degradation 1A (HRD1A), HRD1B, and the Proteins Associated with HRD1-1 (PAWH1 and PAWH2), emerged as the most abundant interactors of PILS2, PILS3, and PILS6 (Fig. 2A). Additionally, while other ERAD components like DERLIN1 (DER1), DER2.1, and DER2.2 were detected in most biological replicates, their presence was not consistent across all samples. This set of data suggests that PILS proteins can associate with key components of the ERAD complex.

**Figure 2:**
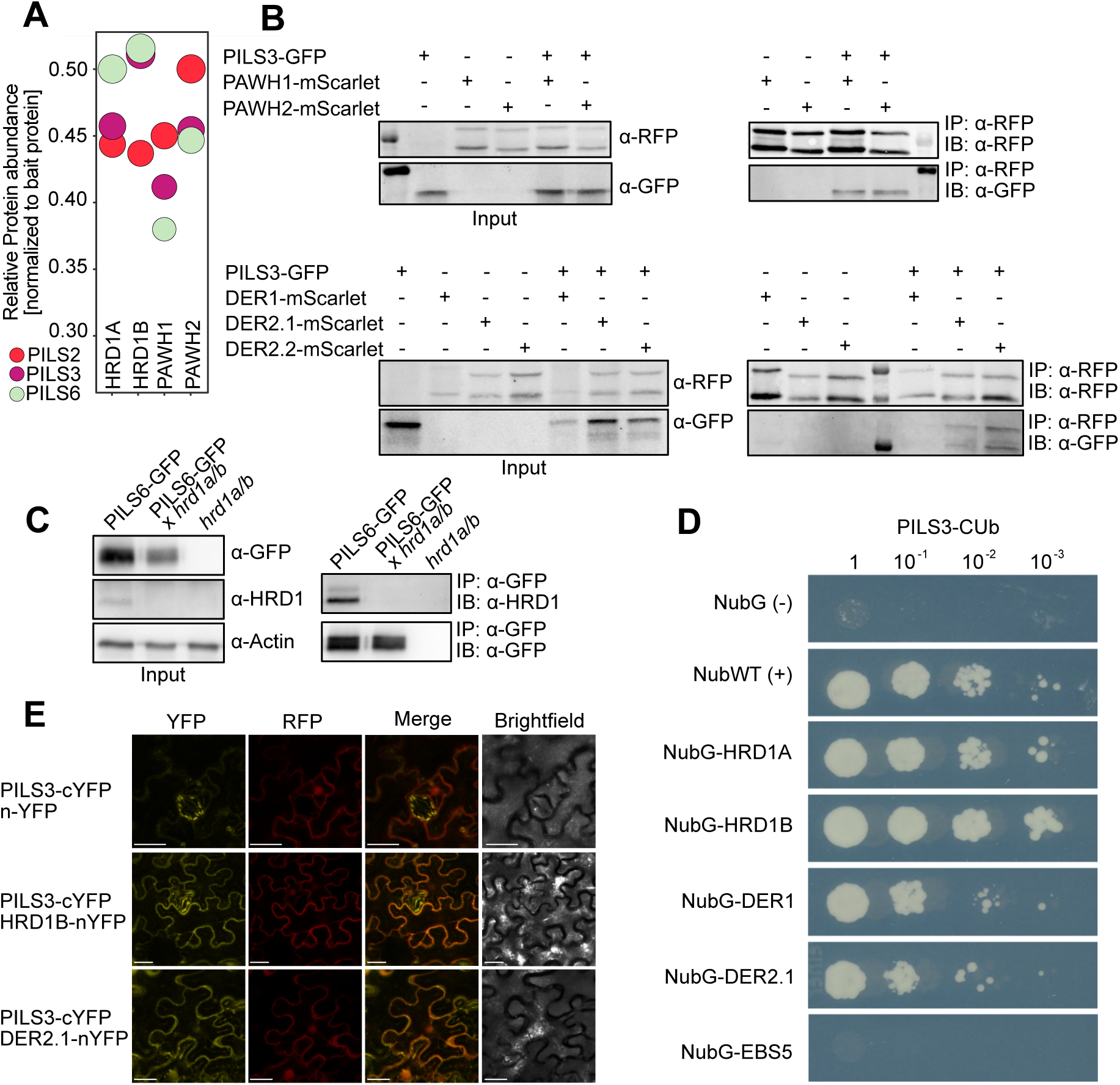
PILS proteins interact with components of the ERAD complex. **A**, Relative protein abundance of co-immunoprecipitated proteins, using GFP-PILS2, GFP-PILS3 and PILS6-GFP as baits. HRD1A/B and PAWH1/2 were among the most abundant interactors from mass spectrometry experiments. **B**, Co-IP of PILS3-GFP with PAWH1/2- mScarlet and DER1/2.1/2.2-mScarlet transiently expressed in *Nicotiana benthamiana*. Inputs and α-RFP or α-GFP immunoprecipitates (IPs) were separated by SDS-PAGE and analysed by immunoblotting (IB) with RFP or GFP antibodies. The same amount of protein was used for each IP. **C**, Co-IP of PILS6-GFP with anti HRD1 antibody in stable *A. thaliana* transgenics. Inputs and α-GFP IPs were separated by SDS-PAGE and analysed by IB with GFP and HRD1 antibodies. The same amount of protein was used for each IP. **D**, Mating-based split-ubiquitin assay between PILS3 and the indicated proteins. Growth of transformed yeast colonies was detected after five days at room temperature. NubG was used as a negative and NubWT as a positive control. **E**, Ratiometric Bimolecular Fluorescence Complementation (rBiFC) in *Nicotiana benthamiana* leaves transiently transformed with constructs encoding PILS3-cYFP and HRD1B-, DER2.1-nYFP, or nYFP alone. A constitutively expressed mRFP (from the same T-DNA) was used as expression control (30). Scale bars, 25 µm. The proteomic screen was performed twice (A), while all other experiments were repeated at least three times.

To further validate our proteomic findings, we co-expressed PILS3-GFP alongside ERAD components, such as PAWH1/2 and DERs, tagged with mScarlet for co-immunoprecipitation (co-IP) in *Nicotiana benthamiana*. Our results demonstrated a strong association between PILS3-GFP and both PAWH1-mScarlet and PAWH2-mScarlet, as well as with DER2.1-mScarlet and DER2.2-mScarlet (Fig. 2B). Using an HRD1 antibody that recognizes endogenous HRD1A/B in *Arabidopsis thaliana*, we further confirmed the interaction of PILS6-GFP with HRD1 in stable transgenic plants (Fig. 2C; fig. S2A). Collectively, these findings support the notion that PILS proteins interact with the ERAD complex.

To investigate whether PILS proteins directly interact with ERAD components, we employed a heterologous system, using a mating-based split-ubiquitin system for transmembrane proteins in yeast (*21*). This analysis suggested direct interactions between PILS3 and HRD1A/B, as well as with DERLIN proteins like DER1 and DER2.1 (Fig. 2D). However, PILS3 did not interact with the regulatory ERAD component EBS5/HRD3, indicating specificity in the observed interactions (Fig. 2D). To consolidate these possibly direct interactions in plant cells, we utilised ratiometric bimolecular fluorescence complementation (rBiFC) (*22*), which allows the normalisation of interaction-dependent signals against a second fluorescent reporter expressed from the same T-DNA. Using this system, we obtained significant interaction signals between PILS3-cYFP and HRD1B-nYFP, DER2.1-nYFP, as well as DER2.2-nYFP when co-expressed in *Nicotiana benthamiana* (Fig. 2E; fig. S2B, C).

To further consolidate this interaction under non-stressed conditions *in planta*, we performed a co-immunoprecipitation using anti-HRD1 antibodies on wild-type *Arabidopsis* lysates, followed by label-free quantitative mass spectrometry. Mock IPs (beads without antibody) were used to assess background binding. This approach confirmed strong enrichment of several canonical ERAD components when compared to control, including HRD1A/B itself as well as EBS5, EBS7, PAWH1 and PAWH2 (fig. S2D). Gene set enrichment analysis of the quantified proteins showed a significant enrichment of proteins related to ERAD and proteolysis. Notably, native PILS5, as well as DER2.1 and DER2.2, were found among the significantly enriched proteins (fig. S2D).

This suggests that PILS proteins can directly interact with ERAD complex components under physiological, non-stressed conditions, categorising them as potential physiological clients of the ERAD system.

The ERAD pathway can target functional proteins, but its canonical role is to identify, extract, and degrade misfolded or aberrant proteins from the ER. We were initially concerned that PILS fusion proteins may be recognised by this quality control mechanism, which may engage with the ERAD pathway. This led us to address whether the PILS/ERAD interactions impose quantitative constraints on the efficient processing of misfolded substrates within this pathway. To explore such a scenario, we utilised the *bri1-5* mutant, which encodes a structurally imperfect yet functional brassinosteroid receptor. In this mutant, BRI1-5 proteins undergo constitutive degradation via the ERAD pathway (*23*). Both pharmacological and genetic interference with ERAD function can reduce the degradation of the BRI1-5 receptor, thereby promoting its secretion to the plasma membrane and partially rescuing the dwarfed phenotype characteristic of this mutant (*23*, *24*).

To quantitatively assess BRI1-5 processing by the ERAD pathway, we performed the well-established Endoglycosidase H (Endo H) assay, which distinguishes the size differences between plasma membrane-localized BRI1 and its ER-localized counterpart based on differences in glycan modifications during transit through the secretory pathway(*23*, *24*). Notably, neither PILS6 overexpression nor loss-of-function mutations in *pils6* affected BRI1-5 processing (fig. S3A), suggesting that BRI1-5 remains retained in the ER. Furthermore, the root phenotype of the *bri1-5* mutant, along with the constitutive degradation of BRI1-5, was not significantly altered in these lines (fig. S3B-D). These findings indicate that overexpression of PILS6-GFP does not interfere with the ERAD-dependent degradation of BRI1-5. Consequently, we conclude that the interaction of PILS6-GFP with ERAD components is unlikely to negatively impact their canonical function.

While the HRD1 branch of the ERAD pathway is well recognised for its role in maintaining ER quality control in plants, it also targets physiological substrates in yeast and animal cells. These physiological ERAD clients are functional proteins that are conditionally degraded to fine-tune their activities (*25*). However, the existence of physiological clients of ERAD complexes in plants are just surfacing(*26–28*), hence remaining an exciting area of research. Moreover, physiological HRD1 clients remain largely unknown in plants (*29*). We found that the *hrd1a hrd1b* double mutants exhibited reduced hypocotyl growth in dark conditions, suggesting that the ERAD complex could play a physiological role in regulating growth beyond its traditional function in ER quality control (Fig. 3A). The PILS proteins redundantly limit organ growth, with the *pils3* single mutants already demonstrating accelerated hypocotyl growth compared to wild-type plants (*2*). The *pils3 hrd1a hrd1b* triple mutant partially alleviated the growth defects observed in the *hrd1a hrd1b* double mutants (Fig. 3A). Moreover, auxin-responsive genes were downregulated in *hrd1a hrd1b* mutant (fig. S3E) and its hypocotyl growth was hypersensitive to the exogenous application of auxin when compared to wild type (Fig. 3B). This set of data suggests deregulated auxin-dependent responses and growth in *hrd1a hrd1b* double mutant. Notably, the hypersensitivity to exogenous auxin could be compensated by *PILS3* overexpression. This set of data reveals that HRD1 contributes to plant growth regulation in non-stressed conditions. While additional growth-related defects from *hrd1a hrd1b* mutations cannot be ruled out, our findings suggest that PILS proteins contribute to HRD1-reliant growth control.

**Figure 3:**
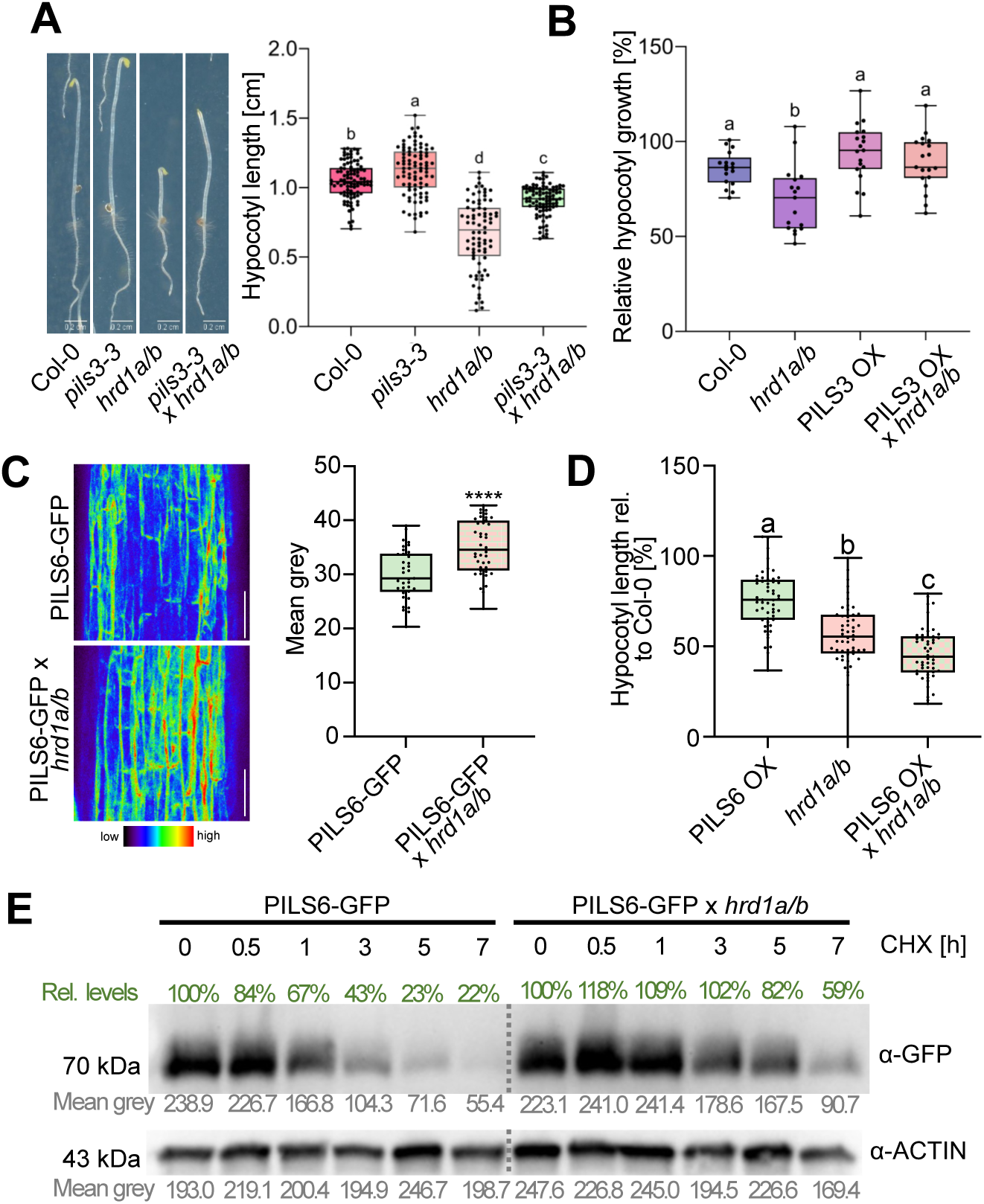
HRD1 activity defines PILS turnover and hypocotyl growth. **A**, Dark-grown hypocotyl length was measured for Col-0 wild type, *pils3-3*, *hrd1a/b*, and *pils3-3 hrd1a/b* mutants. n = 50. one-way ANOVA followed by Tukey’s multiple comparison test (letters indicate statistical differences: P < 0.001). **B**, Relative dark-grown hypocotyl growth (to respective MOCK treatment) of Col-0 wild type, *hrd1a/b*, *p35S::PILS3* overexpressors (PILS3 OX) and PILS3 OX in *hrd1a/b* mutants. Seedlings were germinated on ½ MS plates for 3 days and then transferred to synthetic auxin (NAA 750 nM) or solvent (DMSO) containing plates for 2 days. n = 20. one-way ANOVA followed by Tukey’s multiple comparison test (letters indicate statistical differences: P < 0.001). **C**, Representative images and quantifications of PILS6-GFP and PILS6-GFP x *hrd1a hrd1b* signal in 3-day-old dark-grown seedlings. Scale bars, 50 µm. n = 35-40, Student’s t-test (****P < 0.0001). **D**, Relative hypocotyl length of 5-day-old dark-grown seedlings. n = 45-60, one-way ANOVA followed by Tukey’s multiple comparison test (b, c: P < 0.0001). **E**, Immunoblot analysis of the PILS6-GFP in wild-type or *hrd1a hrd1b* background. 4-day-old seedlings were treated for the indicated time with 100 µM cycloheximide (CHX) in liquid ½ MS, proteins were separated by SDS-PAGE and analysed by immunoblotting using α-GFP antibodies. α-Actin antibody was used for normalization. Values below each band indicate the mean grey value, while the percentage changes relative to the loading control are shown above each band. All panels with boxplots: Box limits represent 25th percentile and 75th percentile; horizontal line represents median. Whiskers display min. to max. values. All experiments were repeated at least three times.

Our previous work on PILS6 proteins revealed a conditional posttranslational control of PILS proteins (*3*). To investigate whether the ERAD complex indeed influences the posttranslational regulation of PILS proteins, we, hence, introduced the constitutive PILS6-GFP expressing lines with the *hrd1a hrd1b* double mutants. The genetic interference with *HRD1* function led to a quantitative increase in PILS6-GFP abundance in dark-grown hypocotyls (Fig. 3C). This suggests that HRD1 is crucial for determining the levels of PILS proteins under non-stressed conditions. Furthermore, the enhanced abundance of PILS6 in the *hrd1a hrd1b* mutant resulted in stronger growth inhibition of dark-grown hypocotyls compared to parental lines (Fig. 3D). These observations suggest that interference with the ERAD pathway stabilises functional PILS proteins, thereby enhancing growth repression.

Next, we used western blotting as a semiquantitative method, employing a cycloheximide (CHX) chase experiment to assess protein turnover. CHX inhibits de novo protein synthesis, allowing us to monitor the degradation dynamics of PILS6-GFP. Following CHX treatment, PILS6-GFP levels steadily declined over time (Fig. 3E). Notably, this decline was significantly attenuated in the *hrd1a hrd1b* double mutant background, indicating that HRD1 activity contributes to the turnover of PILS6-GFP.

To further examine if HRD1 has a direct impact on PILS turnover, we aimed to pharmacologically inhibit HRD1 using LS102, which selectively disrupts HRD1 activity in animal cells (*30*). As mentioned above, the *hrd1a hrd1b* mutants displayed reduced hypocotyl growth in darkness (Fig. 3A, B, D), and this phenotype was phenocopied by LS102 application on wild-type Col-0 plants (fig. S3F). Notably, the LS102-induced effects on hypocotyl expansion were abolished in the *hrd1a hrd1b* double mutants (fig. S3F), indicating that LS102 specifically targets HRD1-dependent processes in dark-grown hypocotyls of Arabidopsis.

Consistent with the genetic interference, pharmacological inhibition of HRD1 significantly increased the protein levels of GFP-PILS3, PILS5-GFP, and PILS6-GFP, which is reminiscent to the inhibition of the proteasome (Fig. 4A, B). Moreover, blocking HRD1 function also resulted within hours in a relatively fast increase in the abundance of functional pPILS3::PILS3-GFP (Fig. 4C). In contrast, PILS3 expression was not affected during this experimental condition (fig. S1E). Collectively, these results suggest that the HRD1-reliant ERAD complex has a direct role in modulating the physiologically relevant levels of PILS proteins.

**Figure 4:**
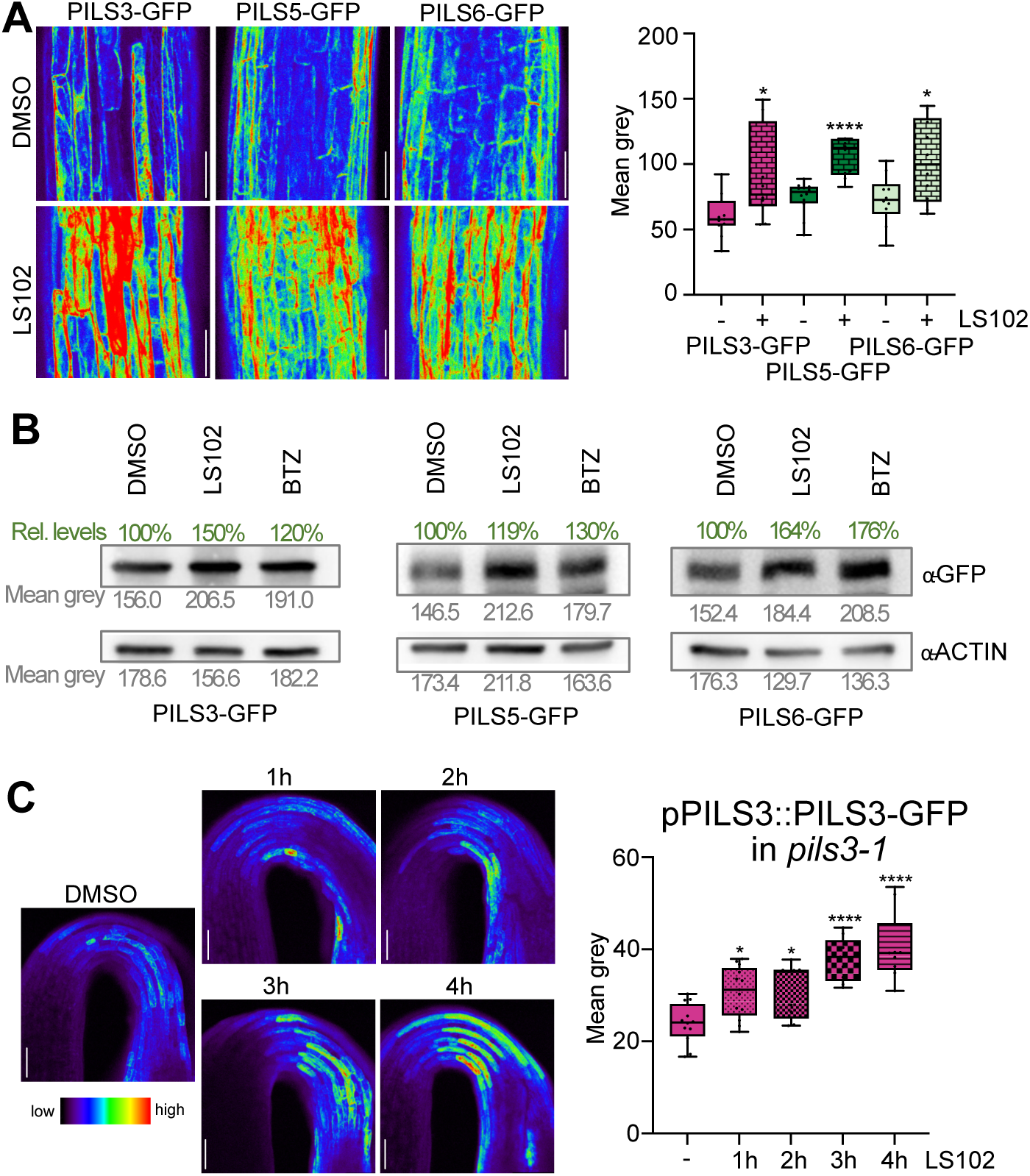
Inhibition of HRD1 has an immediate effect on PILS protein stability **A,** Representative images and quantifications of GFP-PILS3, PILS5-GFP and PILS6-GFP signal in 3-day-old dark-grown seedlings. Seedlings were grown on solid ½ MS and treated with DMSO or 5 µM LS102 (HRD1 inhibitor) in liquid ½ MS for 4h. Scale bars, 50 µm. n = 10-12, Student’s t-test between DMSO and LS102. **B,** Immunoblot of 4-day-old dark-grown GFP-PILS3, PILS5-GFP and PILS6-GFP seedlings treated with DMSO, 10 µM LS102, or 50 µM BTZ for 4 hours in liquid ½ MS. Anti-Actin antibody was used for normalisation. Values below each band indicate the mean grey value (in grey) and the percentage changes relative to the loading control are shown above each band (in green). C, Representative images and quantifications of pPILS3::PILS3-GFP in the *pils3-1* background in 3-day-old dark-grown seedlings. Seedlings were grown on solid ½ MS and treated with DMSO or 5 µM LS102 in liquid ½ MS for 1-4h. n = 10, one-way ANOVA followed by Tukey’s multiple comparison between DMSO and LS102. In all panels with boxplots: Box limits represent 25th percentile and 75th percentile; horizontal line represents median. Whiskers display min. to max. values. P-Values: * P < 0.05, ** P < 0.01, *** P < 0.001, **** P < 0.0001. All experiments were repeated at least three times.

To genetically increase ERAD activity, we turned to EBS5/HRD3, because it plays a conserved role in enhancing HRD1 activity (*24*, *31*). Notably, the introduction of an additional copy of pEBS5:EBS5 leads to the mild upregulation of *EBS5* transcript (fig. S3G). Upregulation of EBS5 correlated with reduced PILS protein levels, such as in the constitutive p35S::PILS5-GFP expression line (fig. S4A). This reduction in PILS5 levels correspondingly balanced the dark-grown hypocotyl phenotype of PILS5 overexpressing seedlings (fig. S4B), suggesting that the degraded PILS5 proteins were functional.

Our findings collectively reveal that the ERAD pathway directly regulates the turnover of functional PILS proteins in non-stressed conditions, ultimately influencing organ growth. To further validate this role, we next explored the requirement of HRD1 in the conditional control of PILS turnover. We previously demonstrated that both internal and external signals, such as the phytohormone brassinosteroid and elevated ambient temperatures, induce the turnover of at least PILS1, PILS2, PILS3, PILS5 and PILS6 proteins in roots, impacting auxin-dependent growth (*3*, *5*).

To investigate the conditional turnover of PILS proteins in the main roots, we applied brassinolide (BL) (fig. S4C) or transferred constitutively PILS6-GFP expressing plants to elevated temperatures (fig. S4D). As expected, both BL treatment and high temperature induced the PILS6 turnover in the wild type. The conditional induction of PILS6 degradation was, however, significantly reduced in *hrd1a hrd1b* mutants (fig. S4C, D), which also affected the respective root organ growth rates ((*3*, *5*); fig. S4E, F). This suggests that the ERAD complex regulates the conditional turnover of functional PILS proteins.

Importantly, we previously revealed that ER-stress responses can modulate PILS proteins in a posttranslational manner (*6*). Notably, both brassinosteroid application and high temperatures intersect with ER stress responses (*32*, *33*) and hence may have a general impact on ER homeostasis. Thus, we next aimed to identify specific conditions under which individual PILS proteins are selectively regulated without globally triggering the ER stress pathway. Given that changes in PILS protein turnover may alleviate root growth repression (*3*, *5*), we utilised root growth dynamics in overexpression lines of PILS2, PILS3, PILS5, and PILS6 to screen for selective modulators of PILS protein turnover.

Considering the previously reported influences of auxin and brassinosteroids on PILS abundance (*5*, *7*), we first tested other phytohormones for a potential impact on PILS protein stability. Interestingly, we found that the application of low concentrations of the phytohormone abscisic acid (ABA) specifically rescued the short root phenotype of PILS3-overexpressing lines, whereas PILS2, PILS5, and PILS6 overexpressors and wild-type roots were largely not affected in these low hormone concentrations (Fig. 5A). In agreement with this observation, these low doses of ABA led to a reduction in the protein abundance of PILS3, while PILS6-GFP showed no detectable changes (Fig. 5B-E). In agreement with a specific effect on PILS3, ER-stress-induced genes remained inactive under these experimental conditions (fig. S5A, B). This set of data reveals that low doses of ABA specifically induce the turnover of PILS3 in an ER-stress-independent manner.

**Figure 5:**
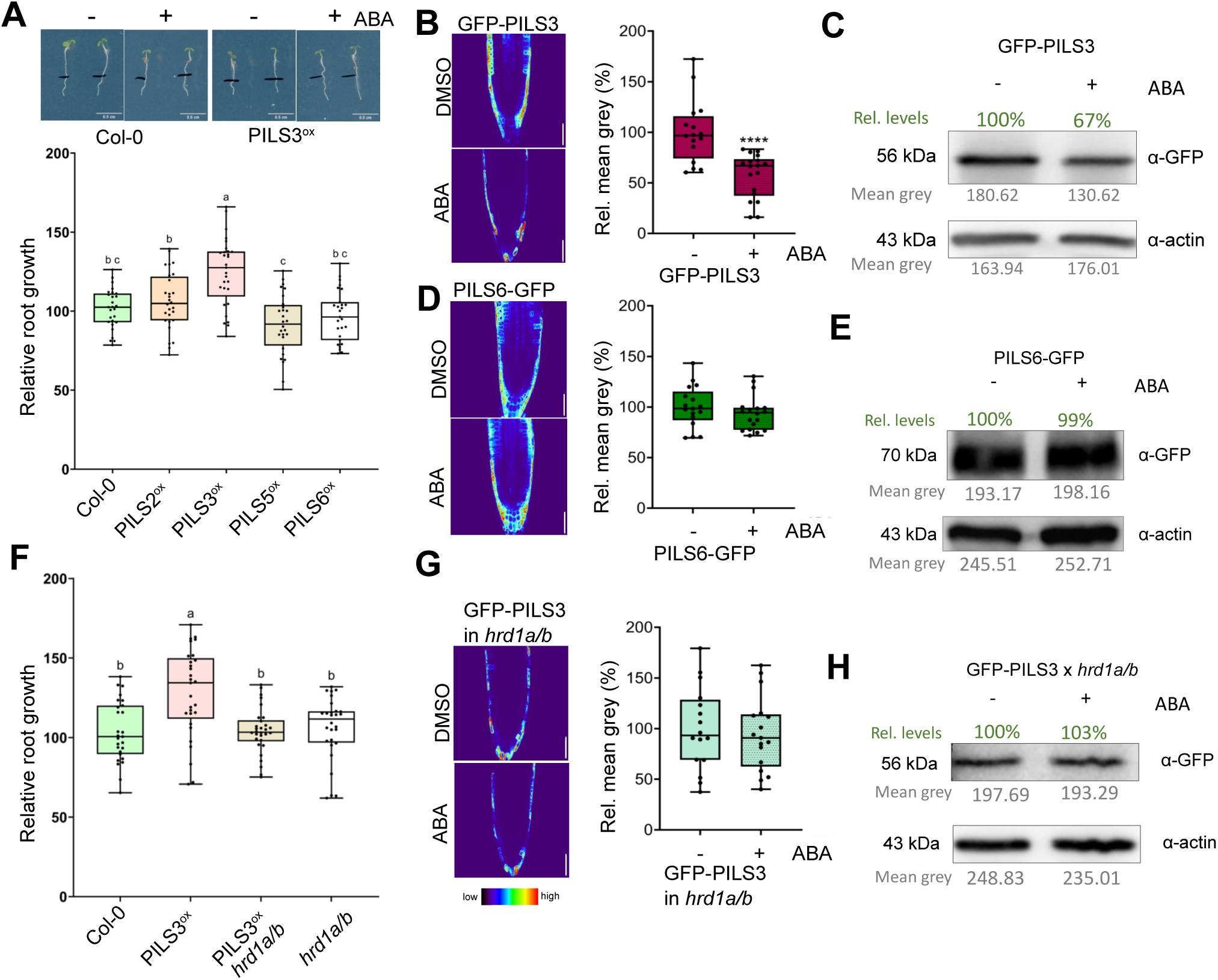
HRD1 regulates conditional PILS3 turnover. **A,** Representative images and relative root growth quantification of 3-day-old seedlings, which were transferred for 1 day on ½ MS media containing DMSO or 100 nM ABA. n = 28-31, one-way ANOVA followed by Tukey’s multiple comparison test (letters indicate statistical differences: P < 0.0001). **BG**, Representative images, its quantifications (B, D, G) and immunoblots (C, E, H) of GFP-PILS3 (B, C), PILS6-GFP (D, E), and GFP-PILS3 in hrd1a/b (G, H) signal in 4-day-old seedlings. Seedlings were grown on solid ½ MS and treated with DMSO or 100 nM ABA in liquid ½ MS for 4h. Scale bars, 50 µm. n = 16-20 from replicates pooled after confirming consistency across experiments, Student’s t-test between DMSO and ABA. Student’s t-test (****P < 0.0001). Proteins were separated by SDS-PAGE and analysed by immunoblotting using α-GFP, and α-Actin antibody was used for normalization, values below each band (in grey) indicate the mean grey value, and the percentage changes relative to the loading control are shown above each band (in green). Seedlings were grown for 4 days on solid ½ MS and treated with DMSO or 100 nM ABA for 4h. **F,** Relative root growth quantification of 3-day-old seedlings, which were transferred for 1 day on ½ MS media containing DMSO or 100 nM ABA. n = 28-31, one-way ANOVA followed by Tukey’s multiple comparison test (a: P < 0.001). In all panels with boxplots: Box limits represent the 25th percentile and 75th percentile; the horizontal line represents the median. Whiskers display min. to max. values. All experiments were repeated at least three times.

Next, we tested if the proteasome and HRD1 are required for the ABA-induced degradation of PILS3. Using initially a pharmacological approach, we revealed that the ABA-induced downregulation of PILS3 was blocked by the inhibition of the proteasome and HRD1 (fig. S6A, B), suggesting an ERAD-dependent regulation. In agreement, the ABA-dependent regulation of PILS3, along with the differential root growth responses in PILS3 overexpressors, was abolished in the *hrd1a hrd1b* mutant background (Fig. 5F-H). Accordingly, we conclude that ABA specifically induces PILS3 degradation in an ERAD-dependent manner. These findings highlight the necessity of the HRD1-reliant ERAD for the selective degradation of PILS proteins.

Collectively, our data indicate that functional PILS proteins are conditionally recruited to the ERAD complex even in the absence of ER stress conditions, initiating their proteasome-dependent turnover. Beyond its established role in ER quality control for plants, we propose that the HRD1-reliant ERAD complex also fulfils a crucial physiological role— regulating the conditional turnover of functional proteins and influencing plant growth regulation.

## Discussion

In our previous work, we demonstrated that ER stress increases and decreases the stability of PILS proteins in a tissue-specific manner (*6*), contributing to auxin-dependent stress adaptation responses. However, the molecular machinery responsible for controlling PILS turnover remained undefined. In the present study, we now identify specific ERAD components—particularly the HRD1 E3 ligases—as interactors of PILS proteins as well as regulators of PILS turnover under non-stress conditions. Endogenous PILS proteins are expressed at low levels, and here we mainly employed functional PILS-GFP fusion constructs to monitor PILS protein dynamics. These reporter lines were previously validated to rescue the respective mutant phenotypes (*1–3*), providing suitable tools for analysing PILS turnover and function *in planta*. We fully recognise the importance of distinguishing artefactual degradation of tagged proteins from bona fide ERAD-mediated turnover of physiological substrates. To ensure the relevance of our findings, we took multiple complementary measures: We initially used constitutive PILS-GFP expression lines to specifically address the posttranslational protein control in the *hrd1a hrd1b* mutant background. We additionally used pharmacological inhibition of proteasome (BTZ) and HRD1 (LS102) activity in functional pPILS3::PILS3-GFP lines, thus confirming these effects under endogenous expression leves. Moreover, we specifically tested whether PILS overexpression leads to unspecific ERAD activation by using the *bri1-5* misfolded receptor allele as a reporter. We observed no evidence for ER stress induction or global ERAD interference, arguing against saturation or non-specific engagement of the ERAD machinery. Our conclusions are further supported by converging independent data where genetic (*hrd1a hrd1b*), pharmacological (LS102), and physiological (ABA, BR, heat) cues all consistently alter PILS turnover in a proteasome-dependent manner and modulate auxin-responsive growth outcomes. Moreover, *hrd1a hrd1b* double mutants are defective in auxin responses and growth, which is genetically modulated by changes in *PILS* activity. Importantly, the selective destabilisation of PILS3 under ABA treatment, while other PILS members remain unaffected, reveals a context-specific ERAD targeting mechanism, not generally attributable to artefacts of GFP fusion proteins. Taken together, our findings provide accumulating functional, genetic, and biochemical evidence that PILS proteins are physiological clients of the HRD1-dependent ERAD pathway, extending its function in plants beyond canonical quality control to include developmentally relevant hormone transport regulation.

The identification of PILS proteins as physiological clients of the ERAD complex opens new avenues for understanding how plants regulate ER protein levels in response to various internal and external stimuli. Notably, HRD1-dependent ERAD pathways have been suggested to selectively target properly folded proteins for degradation in non-plant systems, thereby fine-tuning their physiological levels (*25*). Our findings suggest that HRD1 plays an additional role in plant growth coordination. We propose that this additional role of ERAD has profound implications for understanding how plants adapt to fluctuating physiological conditions. Given that PILS proteins are involved in gating nuclear signalling of auxin—a key hormone influencing growth and development—the regulation of their turnover directly impacts auxin availability and homeostasis (*1–3*). Thus, the ability of the ERAD complex to modulate PILS levels could provide a general mechanism through which plants respond to internal and external cues. While the possibly widespread developmental contributions need to be further explored, our findings highlight that ERAD-mediated turnover of PILS proteins is conditional and can be modulated by internal and environmental signals. Such a dynamic adjustment process can optimise plant growth and development, which becomes increasingly important in the face of environmental challenges. It needs to be addressed to what degree other E3 ligases within the ERAD pathway may recognise PILS as physiological targets. This could potentially provide functional redundancy in ER homeostasis and further enhance plant resilience.

Importantly, the conditional degradation of physiological clients, such as PILS proteins, also raises intriguing questions about substrate recognition by the ERAD complex. While the canonical function of HRD1 is to identify, extract, and degrade misfolded or aberrant proteins from the ER (*34*, *35*), the identification of physiologically relevant clients like PILS suggests that the pathway discerns substrates based on regulatory cues. The specific mechanisms through which PILS proteins are recognised by the ERAD machinery remain to be uncovered but could provide fundamental insights into the activity regulation and/or substrate recognition of ERAD components. We propose that the specific, ABA-dependent regulation of PILS3 could function as a suitable model to study the selectivity of HRD1-reliant ERAD activation, which warrants further investigations.

In conclusion, our work transforms our view on HRD1 and ERAD function in plants, impacting the integration of hormone signalling, stress responses, and ER homeostasis, unlocking new perspectives on plant growth regulation.

## Material and methods

### Plant material and growth conditions

*Arabidopsis thaliana* Col-0 (wild type), p35S::GFP-PILS2 (*1*), p35S::GFP-PILS3 (*2*), p35S::PILS5-GFP (*1*), p35S::PILS6-GFP (*3*), pPILS3::PILS3-GFP in *pils3-1* (*2*), p*ils6-1* (*3*), p35S::DER1-mScarlet (*6*) and *bri1-5* (*36*). Seeds were stratified at 4°C for 2 days in the dark. Seedlings were grown vertically on half-strength Murashige and Skoog medium (1/2 MS salts (Duchefa), pH 5.9, 1% sucrose, and 0.8% agar). Plants were grown under long-day (16 h light/8 h dark) or under dark conditions at 20–22°C.

### Genotyping *hrd1a hrd1b* double mutant

*hrda1* and *hrd1b* single mutants were obtained from NASC (SALK_032914 and SALK_061776), crossed and genotyped using primers listed in Table S 2.

### Chemicals and Treatments

Bortezomib (BTZ) (Santa Cruz), LS102 (Merck), Brefeldin A (BFA) (Merck), VAC1 (15), FM4-64 (Merck), Endo H (Merck), cycloheximide (CHX) (Merck), Abscisic Acid (ABA) (Sigma-Merck), and Tunicamycin (TM) (Santa Cruz) were all dissolved in DMSO (Duchefa). Treatments with BTZ and LS102 were performed on 3-4-day seedlings (transferred to supplemented media) or germinated directly on the respective compound. Treatments with ABA and TM were performed on 3-4-day-old seedlings (treated or transferred to supplemented media).

### DNA constructs

The coding DNA sequence (CDS) of PILS3, DER2.1, DER2.2, PAWH1, and PAWH2 and the genomic sequence of EBS5 were amplified by PCR (primer are listed in Table S 2) from cDNA using Q5 High-Fidelity DNA Polymerase (NEB) and cloned either under the 35S promoter together with mScarlet-I for DER2.1, DER2.2, PAWH1 and PAWH2, with GFP for PILS3 or for EBS5 under its own promoter into pPLV03 using Gibson Assembly (NEB).

For split ubiquitin and BiFC assay CDS of PILS3, HRD1A, HRD1B, DER1, DER2.1, DER2.2 and EBS5 were amplified by PCR (primer are listed in Table S2) from cDNA using Q5 High-Fidelity DNA Polymerase (NEB) and cloned for split ubiquitin assay into pDONR221 and for rBiFC into pDONR221-L1L4/L3L2. Subsequently, the bait (PILS3) and preys were recombined into pMetYC-DEST (*37*) and PNX35-DEST (*38*), respectively or into the rBiFC vector (*39*).

The resulting constructs were transformed into Col-0 plants using the floral dipping method (38) or for transient transformation in tobacco plants.

### Microscopy

Confocal microscopy was done with a Leica SP8 (Leica). Fluorescence signals for GFP (excitation 488 nm, emission peak 509 nm), mScarlet-i (excitation 561 nm, emission peak 607 nm) and YFP (excitation 513 nm, emission peak 527 nm) were detected with a 10x, 20x, or 40x (dry and water immersion, respectively) objective. Z-stacks were recorded with a step size of 840 nm. On average, 24 slices were captured, resulting in an average thickness of approximately 20 µm. Image processing was performed using LAS AF lite software (Leica) or ImageJ.

### Transient Transformation

Transient transformation was performed as described by Loos and Castilho (*40*). Subcellular localization/protein extraction was performed three days after transformation.

### Protein Extraction, Immunoprecipitation (IP) and Immunoblot (IB) Analysis

Seedlings or tobacco leaves were ground to fine powder in liquid nitrogen and solubilized with extraction buffer (25 mM TRIS, pH 7.5, 10 mM MgCl_2_, 15 mM EGTA, 75 mM NaCl, 1 mM DTT, 0.1% Tween20, with freshly added proteinase inhibitor cocktail (Roche) and 40 mM L(+) ascorbic acid for tobacco). After spinning down for 60 min at 4°C with 20.000 rpm supernatant was transferred to a new tube and the protein concentration was assessed using the Bradford method. Protein extracts were either used directly for immunoblot with anti-GFP (Roche #11814460001, 1:1,000), anti-RFP (Chromotek #6g6, 1:1,000), anti-BRI (Agrisera #AS12 1859, 1:1,000), anti-HRD1 (Invitrogen #PA5-110899, 1:1,000) or anti-Actin (Sigma #A0480, 1:10,000) and goat anti-mouse IgG (Jackson ImmunoResearch # 115-036-003, 1:10,000) for detection, or for immunoprecipitation with anti-GFP beads (Chromotek #gtma) or anti-RFP beads (Chromotek #rtma) followed by either immunoblot or mass spectrometry. For the Endo H treatment, protein extracts were combined with 10x Glycoprotein Denaturing Buffer (NEB) and denatured by heating at 100°C for 10 minutes. The extracts were then combined with 10X GlycoBuffer 3 (NEB) and Endo H (NEB) and incubated at 37°C for 1 hour. The reaction was then analyzed *via* immunoblot.

### Mass spectrometry

Beads were transferred to new tubes and resuspended in 40 µL of 2 M urea in 50 mM ammonium bicarbonate (ABC). Disulfide bonds were reduced with 10 mM dithiothreitol for 30 min at room temperature before adding 25 mM iodoacetamide and incubating for another 30 min at room temperature in the dark. Remaining iodoacetamide was quenched by adding 5 mM DTT and the proteins were digested with 150 ng trypsin (Trypsin Gold, Promega) at room temperature for 2 hours. The supernatant was transferred to a new tube, the beads were washed with another 30 µL of 2 M urea in 50 mM ABC and the wash combined with the supernatant. After diluting to 1 M urea with 50 mM ABC, another 150 ng trypsin were added and let digest overnight at 37°C in the dark. Digestion was stopped by adding trifluoroacetic acid (TFA) to a final concentration of 0.5%, and the peptides were desalted using C18 Stagetips (Rappsilber et al., 2007).

Peptides were separated on an Ultimate 3000 RSLC nano-flow chromatography system (Thermo Fisher Scientific), using a pre-column for sample loading (Acclaim PepMap C18, 2 cm × 0.1 mm, 5 μm), and a C18 analytical column (Acclaim PepMap C18, 50 cm × 0.75 mm, 2 μm, both Thermo Fisher Scientific), applying a linear gradient from 2% to 35% solvent B (80% acetonitrile, 0.08% formic acid; solvent A 0.1% formic acid) at a flow rate of 230 nL/min over 120 min respectively 60 min. Eluting peptides were analyzed on a Q Exactive HF-X Orbitrap mass spectrometer coupled to the LC via the Proxeon nanospray source, respectively on an Orbitrap Exploris 480 with the FAIMS Pro interface coupled to the LC via the Nanospray Flex ion source (all Thermo Fisher Scientific). Coated emitter tips were obtained from MSWil (PepSep) and New Objectives.

### RNA extraction, cDNA synthesis and quantitative PCR

RNA was extracted from 4-day-old seedlings if not stated otherwise using the InnuPREP Plant RNA Kit (Analytic Jena) according to the manufacturer’s instructions. Before cDNA synthesis, the RNA samples were treated with InnuPREP DNase I (Analytic Jena). 1 µg of RNA was synthesized using FIREScript RT cDNA synthesis KIT (Solis BioDyne), and qPCR was performed using 2x Takyon for SYBR Assay—no ROX (Eurogentec) following the manufacturer’s instructions on a CFX384 Touch Real-Time PCR Detection System (Bio-Rad). Primers used are listed in Table S2 and the genes were normalized against UBQ5 and EIF4.

### Mass spectrometry data acquisition and analysis

The mass spectrometers were operated in data-dependent acquisition mode (DDA). For the Q Exactive HF-X measurements, survey scans were obtained in a mass range of 380-1650 m/z with lock mass activated, at a resolution of 120k at 200 m/z and an AGC target value of 3E6. The 10 most intense ions were selected with an isolation width of 2.0 m/z, for max. 250 ms at a target value of 1E5 and then fragmented in the HCD cell at 27% collision energy. The spectra were recorded in the Orbitrap at a resolution of 30k. Peptides with a charge of +1 or > +6 were excluded from fragmentation, the peptide match feature was set to preferred, the exclude isotope feature was enabled, and selected precursors were dynamically excluded from repeated sampling for 30 seconds.

Experiments on the Orbitrap Exploris were acquired with FAIMS using 2 CVs (−45 V, −60 V) with 1.5 s cycle time. Full scans were obtained over the mass range of 350-1500 m/z, at a resolution of 60k and an AGC target of 100% with maximum injection time set to auto. Precursors of charge 2+ to 6+ were isolated in a 1 m/z window, with a maximum injection time of 100 ms for an AGC target of 100%, and subsequently fragmented applying a normalized collision energy of 28%. MS2 spectra were acquired in the Orbitrap at 15k resolution. Selected precursors were dynamically excluded from repeated sampling for 20 seconds.

Raw data were processed using the MaxQuant software package (version 1.5.2.9 or 1.6.0.16, Tyanova et al., 2016) and the TAIR10 protein database (Dec 2010, www.arabidopsis.org) respectively the Uniprot Arabidopsis reference proteome (release 2020-01, www.uniprot.org) as well as a database of most common contaminants. The search was performed with full trypsin specificity and a maximum of two missed cleavages. Carbamidomethylation of cysteine residues were set as fixed, oxidation of methionine, phosphorylation of serine, threonine and tyrosine, and N-terminal acetylation as variable modifications. For label-free quantification the “match between runs” feature and the LFQ function were activated - all other parameters were left at default.

Results were filtered at a false discovery rate of 1% at protein and peptide spectrum match level. For downstream analysis protein entries with only 1 razor and unique peptide and less than 2 quantification events were removed and missing LFQ values replaced by a constant.

### Split Ubiquitin

For the split ubiquitin assay, yeast strains THY.AP4 and THY.AP5 were transformed with bait and prey constructs, respectively, using a modified protocol from (*41*). Approximately 100 µl of fresh yeast was scraped from YPD plates and resuspended in 200 µl sterile H_2_O. The resuspended yeast was then centrifuged for 5 minutes at 2000 g, and the supernatant was removed. The yeast was then resuspended in 200 µl yeast transformation buffer (40 % PEG 3350, 200 mM LiAc, 100 mM DTT), added 10 µl single-stranded carrier DNA and 1 µg of plasmid DNA and mixed by pipetting up and down. Yeast was incubated for 15 min at 30 °C and for 45 min at 45 °C. The cells were then plated on SD medium and incubated for 4 days at 28 °C. A pool of transformed colonies was mated as described in (*37*). The selected diploid colonies were then incubated on plates containing selective medium (SD -Trp, -Leu, -Ade, - His, -Ura) at 21°C. Growth was recorded up to 9 days after plating.

## Statistical analysis and reproducibility

GraphPad Prism software 9 was used to evaluate the statistical significance of the differences observed between control and treated groups and to generate the graphs. All experiments were, if not stated different, always repeated at least three times and the depicted data show the results from one representative experiment.

## Acknowledgments

We are grateful to Ykä Helariutta for sharing published material; our team members for helpful discussions; to the Max Perutz Labs Mass Spectrometry Facility, LIC Imaging Center Freiburg and Bettina Knapp (Freiburg) for expertise and support. This work was supported by Vienna Research Group (VRG) program of the Vienna Science and Technology Fund (WWTF to J.K-V.), the Austrian Science Fund (FWF) (P29754 to J.K-V., P33497 to S.W. and Hertha Firnberg T728-B16 and Elise Richter V690-B25 to E.F.), the European Research Council (ERC) (639478-AuxinER and 101166880-STARMORPH to J.K-V.), German Science Fund (DFG; 470007283, 559042842, 499026372 to J.K-V. and CIBSS – EXC-2189 to J.K.-V. and P.F.H.) and Austrian Academy of Sciences (25479 to J.F.).

## Author contributions

S.W., J.F., S.N., E.F., M.I.F, L.M.-M., A.-K.R., and D.S. performed experiments. S.N.W.H and P.F.H. contributed the co-IP-MS analysis. R.S. provided the *hrd1a hrd1b* double mutant. S.W. and J.K.-V. devised and coordinated the project and wrote the manuscript. All authors saw and commented on the manuscript.

## Conflict of interest

The authors declare no competing interests.

## Notes

### Competing Interest Statement

The authors have declared no competing interest.

## References

1. E. Barbez, M. Kubeš, J. Rolčík, C. Béziat, A. Pěnčík, B. Wang, M. R. Rosquete, J. Zhu, P. I. Dobrev, Y. Lee, E. Zažímalovà, J. Petrášek, M. Geisler, J. Friml, J. Kleine-Vehn, A novel putative auxin carrier family regulates intracellular auxin homeostasis in plants. Nature 485, 119–122 (2012).

2. C. Béziat, E. Barbez, M. I. Feraru, D. Lucyshyn, J. Kleine-Vehn, Light triggers PILS-dependent reduction in nuclear auxin signalling for growth transition. Nat Plants 3, 17105 (2017).

3. E. Feraru, M. I. Feraru, E. Barbez, S. Waidmann, L. Sun, A. Gaidora, J. Kleine-Vehn, PILS6 is a temperature-sensitive regulator of nuclear auxin input and organ growth in Arabidopsis thaliana. Proc Natl Acad Sci U S A 116, 3893–3898 (2019).

4. J. Mravec, P. Skůpa, A. Bailly, K. Hoyerová, P. Krecek, A. Bielach, J. Petrásek, J. Zhang, V. Gaykova, Y.-D. Stierhof, P. I. Dobrev, K. Schwarzerová, J. Rolcík, D. Seifertová, C. Luschnig, E. Benková, E. Zazímalová, M. Geisler, J. Friml, Subcellular homeostasis of phytohormone auxin is mediated by the ER-localized PIN5 transporter. Nature 459, 1136–1140 (2009).

5. L. Sun, E. Feraru, M. I. Feraru, S. Waidmann, W. Wang, G. Passaia, Z.-Y. Wang, K. Wabnik, J. Kleine-Vehn, PIN-LIKES Coordinate Brassinosteroid Signaling with Nuclear Auxin Input in Arabidopsis thaliana. Curr Biol 30, 1579–1588.e6 (2020).

6. S. Waidmann, C. Béziat, J. Ferreira Da Silva Santos, E. Feraru, M. I. Feraru, L. Sun, S. Noura, Y. Boutté, J. Kleine-Vehn, Endoplasmic reticulum stress controls PIN-LIKES abundance and thereby growth adaptation. Proc Natl Acad Sci U S A 120, e2218865120 (2023).

7. E. Feraru, M. I. Feraru, J. Moulinier-Anzola, M. Schwihla, J. Ferreira Da Silva Santos, L. Sun, S. Waidmann, B. Korbei, J. Kleine-Vehn, PILS proteins provide a homeostatic feedback on auxin signaling output. Development 149, dev200929 (2022).

8. R. Strasser, Protein Quality Control in the Endoplasmic Reticulum of Plants. Annu Rev Plant Biol 69, 147–172 (2018).

9. J. Pollier, T. Moses, M. González-Guzmán, N. De Geyter, S. Lippens, R. Vanden Bossche, P. Marhavý, A. Kremer, K. Morreel, C. J. Guérin, A. Tava, W. Oleszek, J. M. Thevelein, N. Campos, S. Goormachtig, A. Goossens, The protein quality control system manages plant defence compound synthesis. Nature 504, 148–152 (2013).

10. R. D. Etherington, M. Bailey, J.-B. Boyer, L. Armbruster, X. Cao, J. C. Coates, T. Meinnel, M. Wirtz, C. Giglione, D. J. Gibbs, Nt-acetylation-independent turnover of SQUALENE EPOXIDASE 1 by Arabidopsis DOA10-like E3 ligases. Plant Physiology 193, 2086–2104 (2023).

11. Q. Chen, R. Liu, Y. Wu, S. Wei, Q. Wang, Y. Zheng, R. Xia, X. Shang, F. Yu, X. Yang, L. Liu, X. Huang, Y. Wang, Q. Xie, ERAD-related E2 and E3 enzymes modulate the drought response by regulating the stability of PIP2 aquaporins. The Plant Cell 33, 2883–2898 (2021).

12. V. G. Doblas, V. Amorim-Silva, D. Posé, A. Rosado, A. Esteban, M. Arró, H. Azevedo, A. Bombarely, O. Borsani, V. Valpuesta, A. Ferrer, R. M. Tavares, M. A. Botella, The SUD1 Gene Encodes a Putative E3 Ubiquitin Ligase and Is a Positive Regulator of 3-Hydroxy-3-Methylglutaryl Coenzyme A Reductase Activity in Arabidopsis. The Plant Cell 25, 728– 743 (2013).

13. P. Baster, S. Robert, J. Kleine-Vehn, S. Vanneste, U. Kania, W. Grunewald, B. De Rybel, T. Beeckman, J. Friml, SCF(TIR1/AFB)-auxin signalling regulates PIN vacuolar trafficking and auxin fluxes during root gravitropism. EMBO J 32, 260–274 (2013).

14. J. Kleine-Vehn, J. Leitner, M. Zwiewka, M. Sauer, L. Abas, C. Luschnig, J. Friml, Differential degradation of PIN2 auxin efflux carrier by retromer-dependent vacuolar targeting. Proc Natl Acad Sci U S A 105, 17812–17817 (2008).

15. J. Leitner, K. Retzer, B. Korbei, C. Luschnig, Dynamics in PIN2 auxin carrier ubiquitylation in gravity-responding Arabidopsis roots. Plant Signal Behav 7, 1271– 1273 (2012).

16. S. Niemes, M. Labs, D. Scheuring, F. Krueger, M. Langhans, B. Jesenofsky, D. G. Robinson, P. Pimpl, Sorting of plant vacuolar proteins is initiated in the ER. Plant J 62, 601–614 (2010).

17. J. Kleine-Vehn, P. Dhonukshe, M. Sauer, P. B. Brewer, J. Wiśniewska, T. Paciorek, E. Benková, J. Friml, ARF GEF-dependent transcytosis and polar delivery of PIN auxin carriers in Arabidopsis. Curr Biol 18, 526–531 (2008).

18. K. Dünser, M. Schöller, A.-K. Rößling, C. Löfke, N. Xiao, B. Pařízková, S. Melnik, M. Rodriguez-Franco, E. Stöger, O. Novák, J. Kleine-Vehn, Endocytic trafficking promotes vacuolar enlargements for fast cell expansion rates in plants. Elife 11, e75945 (2022).

19. A. F. Kisselev, W. A. van der Linden, H. S. Overkleeft, Proteasome inhibitors: an expanding army attacking a unique target. Chem Biol 19, 99–115 (2012).

20. S. Waidmann, L. De-Araujo, J. Kleine-Vehn, B. Korbei, Immunoprecipitation of Membrane Proteins from Arabidopsis thaliana Root Tissue. Methods Mol Biol 1761, 209–220 (2018).

21. C. Grefen, P. Obrdlik, K. Harter, The determination of protein-protein interactions by the mating-based split-ubiquitin system (mbSUS). Methods Mol Biol 479, 217–233 (2009).

22. D. G. Mehlhorn, N. Wallmeroth, K. W. Berendzen, C. Grefen, 2in1 Vectors Improve In Planta BiFC and FRET Analyses. Methods Mol Biol 1691, 139–158 (2018).

23. Z. Hong, H. Jin, T. Tzfira, J. Li, Multiple mechanism-mediated retention of a defective brassinosteroid receptor in the endoplasmic reticulum of Arabidopsis. Plant Cell 20, 3418–3429 (2008).

24. W. Su, Y. Liu, Y. Xia, Z. Hong, J. Li, Conserved endoplasmic reticulum-associated degradation system to eliminate mutated receptor-like kinases in Arabidopsis. Proc Natl Acad Sci U S A 108, 870–875 (2011).

25. Y. Jo, R. A. DeBose-Boyd, Post-Translational Regulation of HMG CoA Reductase. Cold Spring Harb Perspect Biol 14, a041253 (2022).

26. T. Guo, H. Weber, M. C. E. Niemann, L. Theisl, G. Leonte, O. Novák, T. Werner, Arabidopsis HIPP proteins regulate endoplasmic reticulum-associated degradation of CKX proteins and cytokinin responses. Molecular Plant 14, 1918–1934 (2021).

27. J. Li, B. Zhang, P. Duan, L. Yan, H. Yu, L. Zhang, N. Li, L. Zheng, T. Chai, R. Xu, Y. Li, An endoplasmic reticulum-associated degradation–related E2–E3 enzyme pair controls grain size and weight through the brassinosteroid signaling pathway in rice. The Plant Cell 35, 1076–1091 (2023).

28. S. Tang, Z. Zhao, X. Liu, Y. Sui, D. Zhang, H. Zhi, Y. Gao, H. Zhang, L. Zhang, Y. Wang, M. Zhao, D. Li, K. Wang, Q. He, R. Zhang, W. Zhang, G. Jia, W. Tang, X. Ye, C. Wu, X. Diao, An E2-E3 pair contributes to seed size control in grain crops. Nat Commun 14, 3091 (2023).

29. G. Langin, M. Raffeiner, D. Biermann, M. Franz-Wachtel, D. Spinti, F. Börnke, B. Macek, S. Üstün, ER-anchored protein sorting controls the fate of two proteasome activators for intracellular organelle communication during proteotoxic stress. [Preprint] (2023). 10.1101/2023.12.11.571118.

30. N. Yagishita, S. Aratani, C. Leach, T. Amano, Y. Yamano, K. Nakatani, K. Nishioka, T. Nakajima, RING-finger type E3 ubiquitin ligase inhibitors as novel candidates for the treatment of rheumatoid arthritis. Int J Mol Med 30, 1281–1286 (2012).

31. N. Vashistha, S. E. Neal, A. Singh, S. M. Carroll, R. Y. Hampton, Direct and essential function for Hrd3 in ER-associated degradation. Proc Natl Acad Sci U S A 113, 5934– 5939 (2016).

32. P. Che, J. D. Bussell, W. Zhou, G. M. Estavillo, B. J. Pogson, S. M. Smith, Signaling from the endoplasmic reticulum activates brassinosteroid signaling and promotes acclimation to stress in Arabidopsis. Sci Signal 3, ra69 (2010).

33. E. M. Neill, M. C. R. Byrd, T. Billman, F. Brandizzi, A. E. Stapleton, Plant growth regulators interact with elevated temperature to alter heat stress signaling via the Unfolded Protein Response in maize. Sci Rep 9, 10392 (2019).

34. J. Schoberer, U. Vavra, Y.-J. Shin, C. Grünwald-Gruber, R. Strasser, Elucidation of the late steps in the glycan-dependent ERAD of soluble misfolded glycoproteins. Plant J 121, e17185 (2025).

35. K. Wu, S. Itskanov, D. L. Lynch, Y. Chen, A. Turner, J. C. Gumbart, E. Park, Substrate recognition mechanism of the endoplasmic reticulum-associated ubiquitin ligase Doa10. Nat Commun 15, 2182 (2024).

36. T. Noguchi, S. Fujioka, S. Choe, S. Takatsuto, S. Yoshida, H. Yuan, K. A. Feldmann, F. E. Tax, Brassinosteroid-insensitive dwarf mutants of Arabidopsis accumulate brassinosteroids. Plant Physiol 121, 743–752 (1999).

37. C. Grefen, S. Lalonde, P. Obrdlik, Split-ubiquitin system for identifying protein-protein interactions in membrane and full-length proteins. Curr Protoc Neurosci Chapter 5, Unit 5.27 (2007).

38. C. Grefen, M. R. Blatt, A 2in1 cloning system enables ratiometric bimolecular fluorescence complementation (rBiFC). Biotechniques 53, 311–314 (2012).

39. A. Hecker, N. Wallmeroth, S. Peter, M. R. Blatt, K. Harter, C. Grefen, Binary 2in1 Vectors Improve in Planta (Co)localization and Dynamic Protein Interaction Studies. Plant Physiol 168, 776–787 (2015).

40. A. Castilho, Ed., Glyco-Engineering: Methods and Protocols (Springer New York, New York, NY, 2015; https://link.springer.com/10.1007/978-1-4939-2760-9)vol. 1321 of Methods in Molecular Biology.

41. R. D. Gietz, R. A. Woods, Transformation of yeast by lithium acetate/single-stranded carrier DNA/polyethylene glycol method. Methods Enzymol 350, 87–96 (2002).

