## Supplemental figures 1-6 for "ERAD machinery controls the conditional turnover of PIN-LIKES in plants"

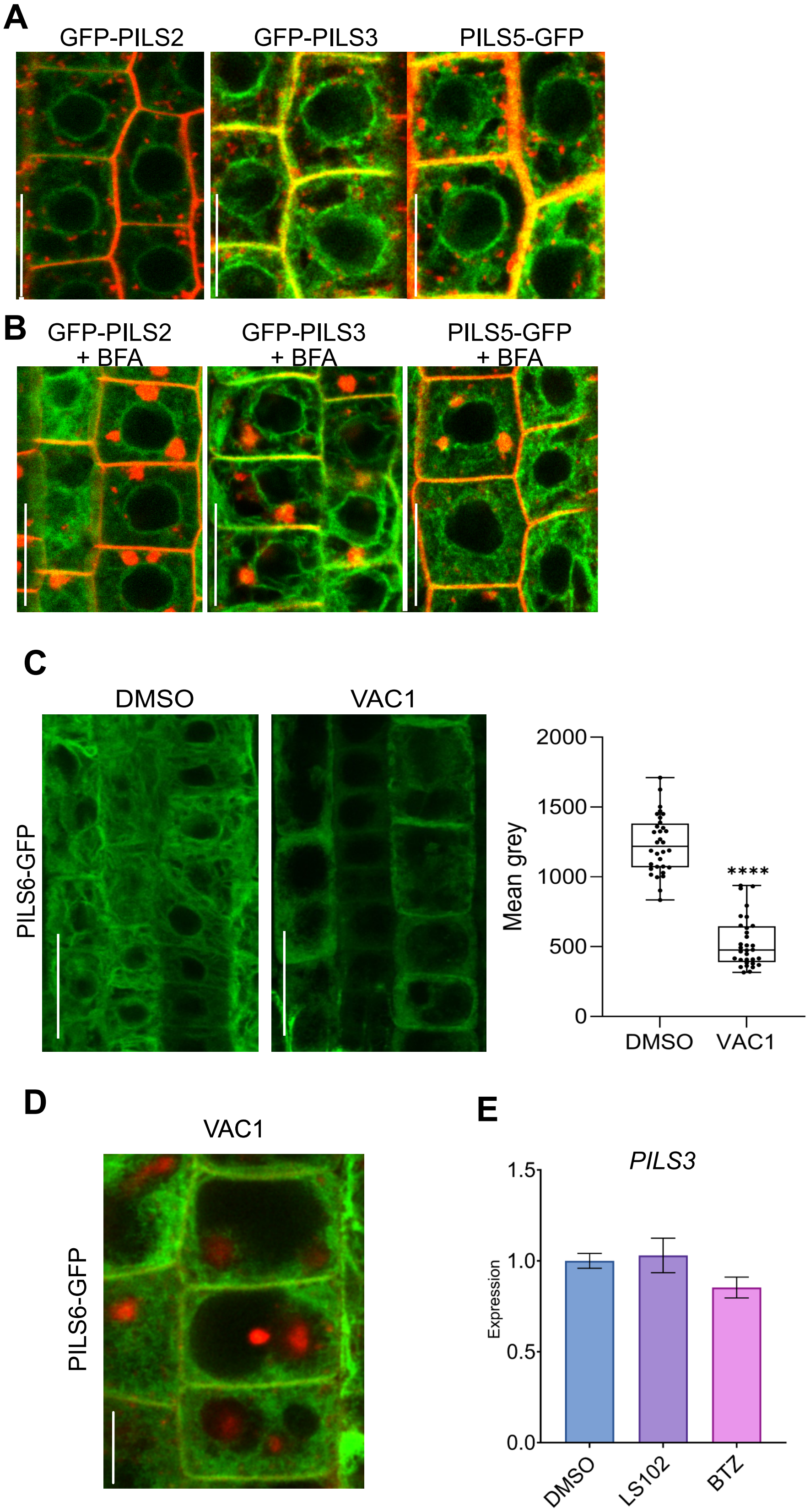


**Figure S1: PILS proteins are stably retained in the ER.**
**A-B,** Representative images of untreated (A) and Brefeldin A (BFA)-treated (50 µM for 2h) (B) GFP-PILS2, GFP-PILS3 and PILS5-GFP. The endocytic dye FM4-64 was used as a counterstain and to illustrate BFA-induced endomembrane accumulations. Scale bars, 25 µm. **C,** Representative images and quantification of PILS6-GFP signal in roots, which were treated with solvent control (DMSO) or 10 µM Vacuolar Affecting Compound 1 (VAC1) for 1h. Box limits represent the 25th percentile and 75th percentile; the horizontal line represents the median. Whiskers display min. to max. values. n > 26, Student's t-test (****P < 0.0001). **D,** PILS6 was absent from VAC1-induced accumulations as indicated by FM4-64 staining. Scale bars, 10 µm. Experiments were done in liquid ½ MS medium using 5-day-old seedlings. All experiments were repeated at least three times. **E,** qPCR analysis of GFP transcript levels in pPILS3::PILS3-GFP expressed in *pils3-1* background. Transcript levels were normalised against *UBQ5* and *EIF4* in 4-day-old dark-grown hypocotyls after being treated with 5 µM LS102 or 40 µM BTZ for 1 hour. Bars represent means ± SD, n = 3.

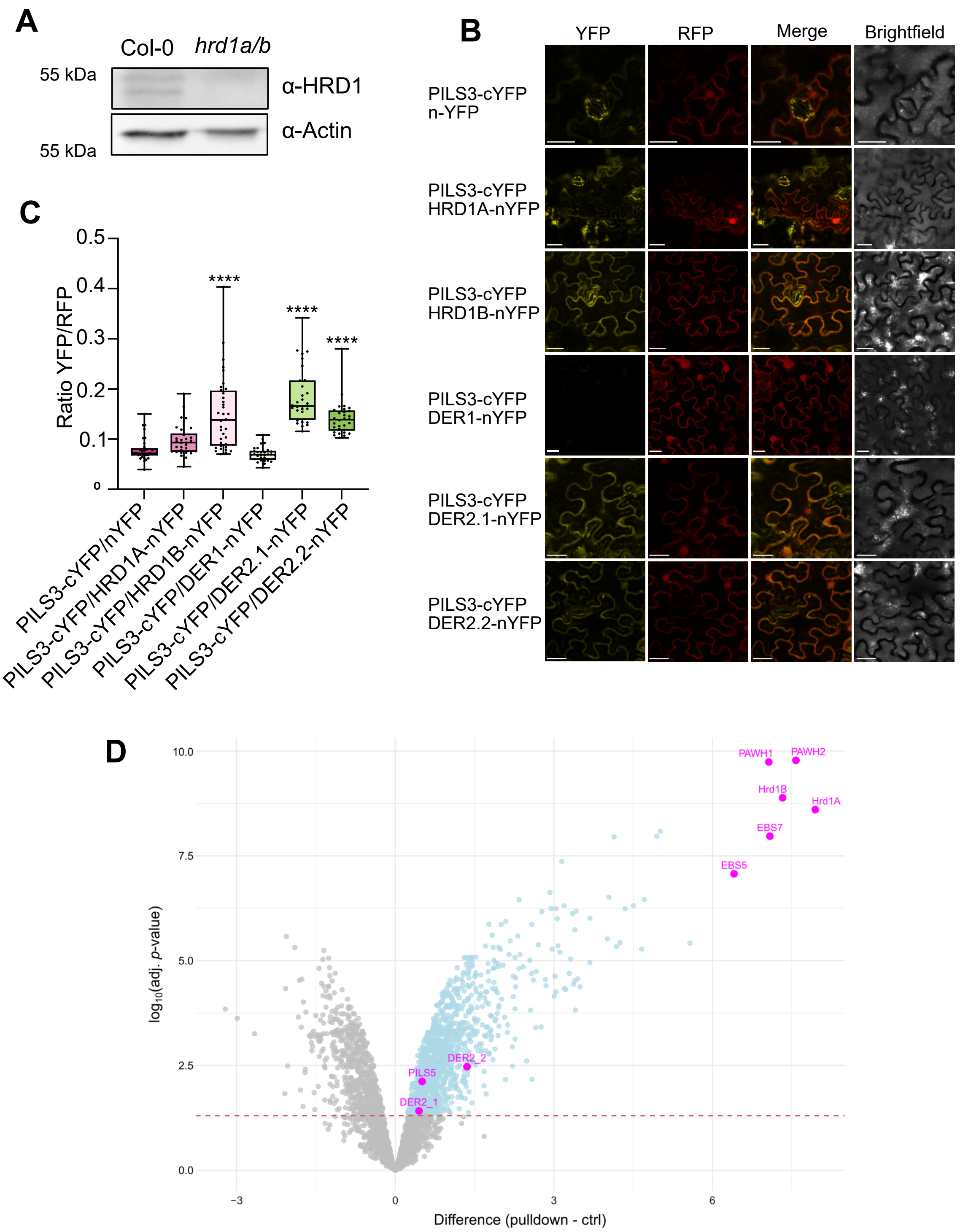


**Figure S2: PILS3 directly interacts with ERAD complex components.**

**A,** Immunoblot of 7-days-old Col-0 and hrd1a hrd1b seedlings. Proteins were separated by SDS-PAGE and analysed by immunoblotting using α-HRD1 antibodies. α-Actin antibody was used for normalization. **B,** Ratiometric BioFluorescence Complementation (rBiFC) in Nicotiana benthamiana leaves transiently transformed with constructs encoding PILS3-cYFP and HRD1A-, HRD1B-, DER1-, DER2.1-, DER2.2-nYFP, or nYFP alone. A constitutively expressed mRFP (from the same T-DNA) was used as expression control (30). Scale bars, 25 µm. **C,** Quantification of BiFC signal was achieved by calculating the ratio between complemented YFP to RFP from 6 entire individual images and from 5 individual interfaces of two to three cells to avoid bias arising from picking single cells. One-way ANOVA followed by Tukey's multiple comparison test (****P < 0.0001). Box limits represent the 25th percentile and 75th percentile; the horizontal line represents the median. Whiskers display min. to max. values. All experiments were repeated at least three times. **D,** Volcano plot of IP-MS analysis showing the difference between log2 values of normalized label-free quantitation intensity values (LFQ) as a measure of protein abundance, plotted against the negative log10 of the adjusted limma-moderated p-value. Blue, proteins significantly co-enriched with Hrd1 (limma-moderated p-value < 0.05 after adjustment for multiple hypothesis testing), indicating potential interactors. Hrd1 and selected proteins of interest are depicted in magenta.

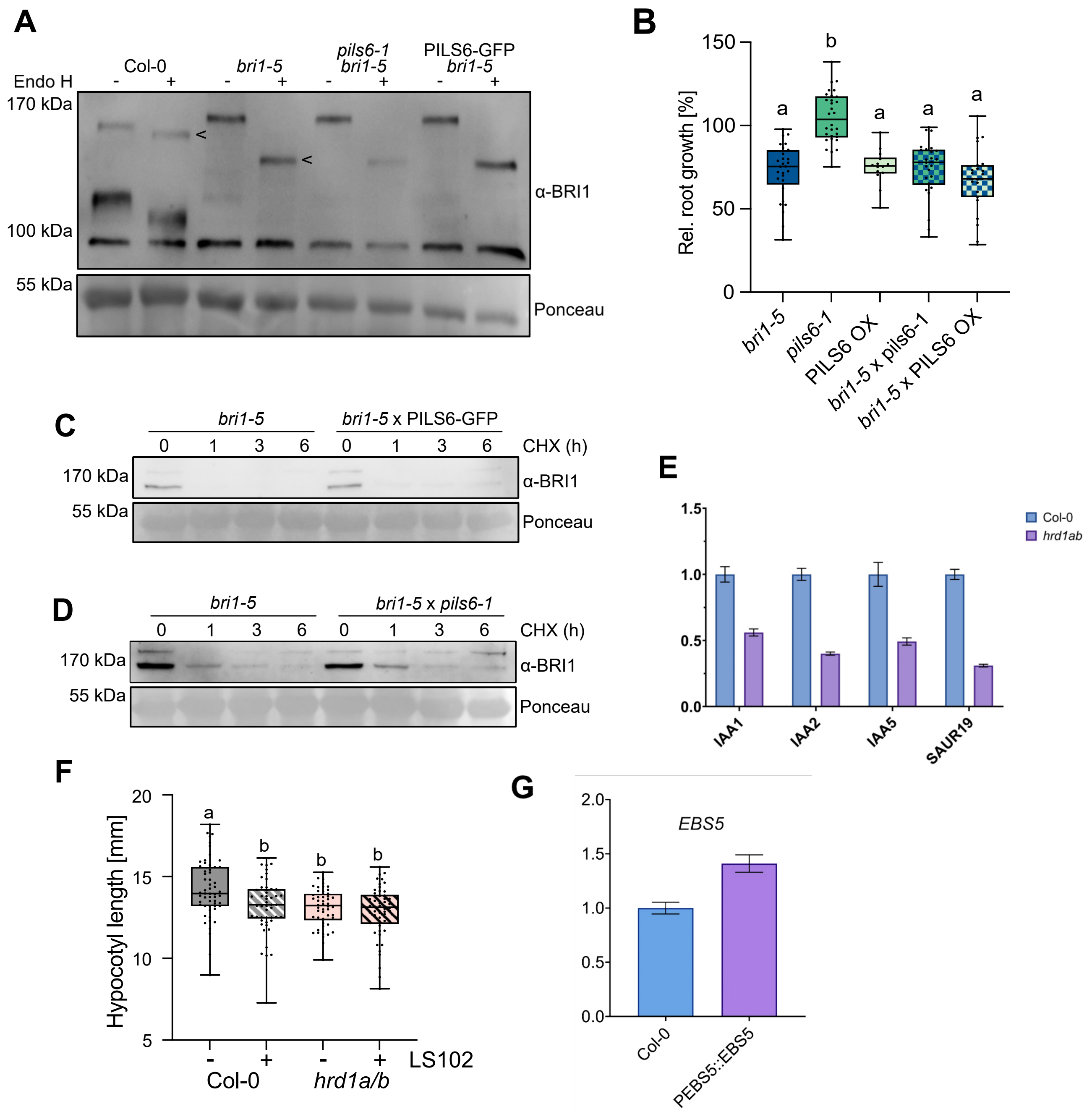


**Figure S3: PILS does not affect ERAD-dependent BRI1-5 processing.**
**A,** Immunoblot of protein extracts from 7-day-old seedlings, treated with (+) or without (-) Endo H for 1 hour. Proteins were separated by SDS-PAGE and analysed by immunoblotting using α-BRI1 antibodies. Ponceau staining was used for normalization. Arrowheads pinpoint size difference of PM and ER localised BRI1 receptors. **B,** Relative root length of 5-day-old seedlings. n = 15-30, One-way ANOVA followed by Tukey's multiple comparison test (b: $P < 0.001$). Box limits represent 25th percentile and 75th percentile; the horizontal line represents the median. Whiskers display min. to max. values. **C, D,** Immunoblot from 5-day-old seedlings treated for the indicated time with 100 µM cycloheximide (CHX) in liquid ½ MS. Proteins were separated by SDS-PAGE and analysed by immunoblotting using α-BRI1 antibodies. Ponceau staining was used for normalization. **E,** qPCR analysis of auxin auxin-responsive genes detecting transcript levels of *IAA1*, *IAA2*, *IAA5*, and *SAUR19* in Col-0 and *hrd1a/b* mutants. Transcript levels were normalised against *UBQ5* and *EIF4*. 4-day-old dark-grown seedlings were used for RNA extraction. Bars represent means ± SD, n = 3. **F,** Hypocotyl length of 2-day-old dark-grown seedlings treated for 24h with DMSO or 10 µM LS102 in liquid ½ MS media. n = 50. one-way ANOVA followed by Tukey's multiple comparison test (b: $P < 0.001$). All experiments were repeated at least three times. **G,** qPCR analysis of *EBS5* transcript levels in Col-0 and pEBS5::EBS5 expressing transgenics. Transcript levels were normalised against *UBQ5* and *EIF4* in 2-week-old seedlings. Bars represent means ± SD, n = 3.

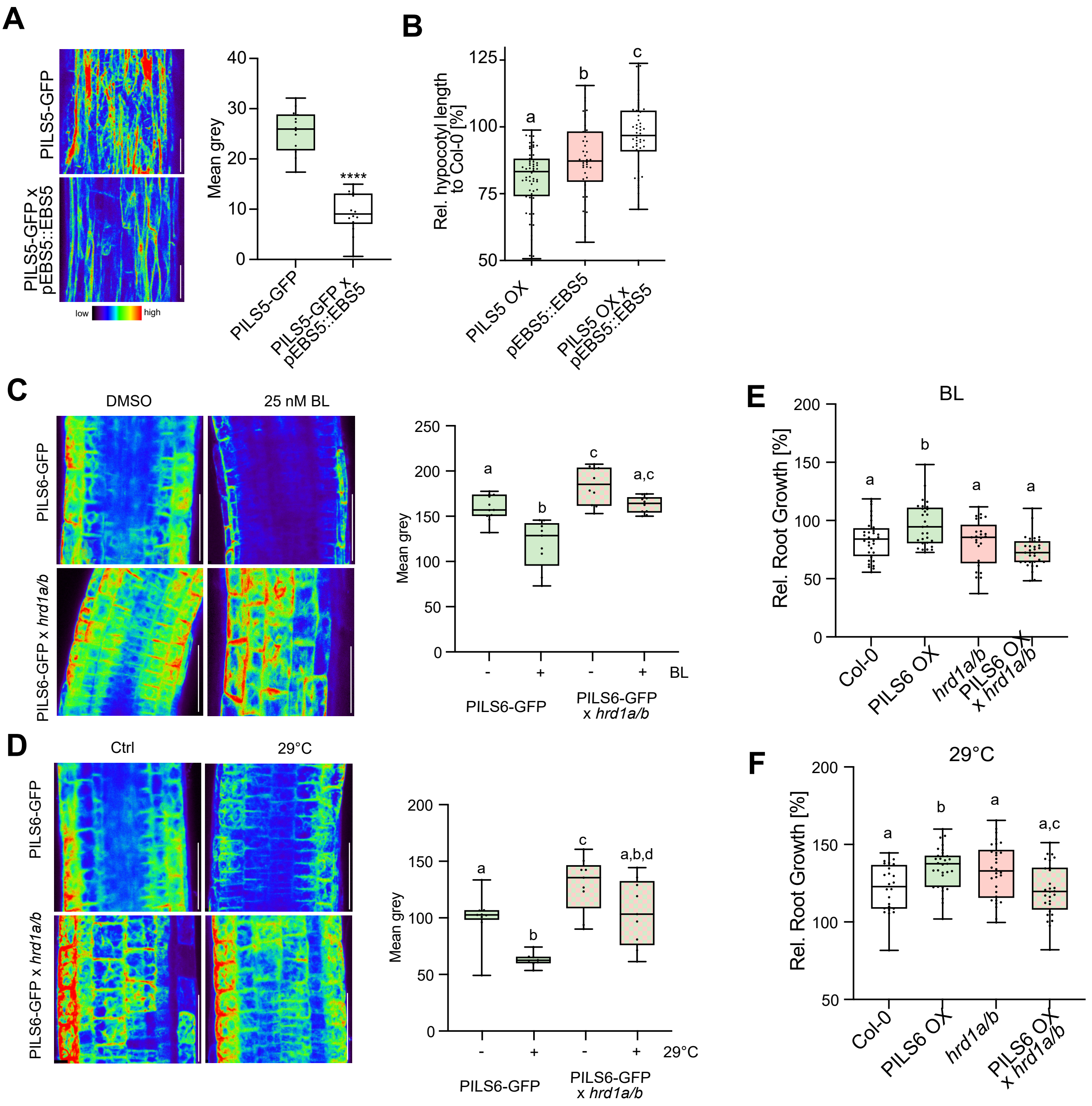


**Figure S4: HRD1 integrates internal and external signals with PILS turnover.**

**A,** Representative images and quantifications of PILS5-GFP and PILS5-GFP x pEBS5::EBS5 signal in 3-days-old dark-grown seedlings. Scale bars, 50 µm. n = 14, Student's t-test (****P < 0.0001). **B,** Relative hypocotyl length of 3-day-old dark-grown seedlings. n = 45-60. one-way ANOVA followed by Tukey's multiple comparison test (b: pEBS5::EBS5 vs. Col-0 P < 0.05, pEBS5::EBS5 vs. PILS5 OX x pEBS5::EBS5 P < 0.01; c: PILS5 OX x pEBS5::EBS5 vs. PILS5 OX P < 0.001). **C, D,** Representative images and quantifications of PILS6-GFP and PILS6-GFP x *hrd1a hrd1b* signal in roots. 3-days-old seedlings were treated (C) for 2h in liquid ½ MS with DMSO or 25 nM 24-Eppibrassinolide (BL) or **(D)** plates were transferred for 72h to 29°C or kept as control at 21°C. Scale bars, 50 µm. n = 9-11, two-way ANOVA followed by Tukey's multiple comparison test (**A,** b: PILS6-GFP ctrl. vs. BL and PILS6-GFP BL vs. PILS6-GFP x *hrd1a hrd1b* ctrl. P < 0.001, PILS6-GFP BL vs. PILS6-GFP x *hrd1a hrd1b* BL. P < 0.0001; c: PILS6-GFP ctrl. vs. PILS6-GFP x *hrd1a hrd1b* ctrl. P < 0.05; B, b: PILS6-GFP ctrl. vs. 29°C and PILS6-GFP 29°C vs. PILS6-GFP x *hrd1a hrd1b* 29°C. P < 0.01, PILS6-GFP 29°C vs. PILS6-GFP x *hrd1a hrd1b* ctrl. P < 0.001; c: PILS6-GFP ctrl. vs. PILS6-GFP x *hrd1a hrd1b* ctrl. P < 0.05. **E, F,** Relative root length of 3-day-old seedlings which were transferred for 2d to ½ MS media containing DMSO or 100 nM BL **(E)** or 4-day-seedlings which were transferred for 3d to 29°C or kept at 21°C (F). n = 25-33, two-way ANOVA followed by Tukey's multiple comparison test (*E,* b: Col-0 vs. PILS6 OX P < 0.05; PILS6 OX vs. *hrd1a hrd1b* P < 0.01; PILS6 OX vs. PILS6 OX x *hrd1a hrd1b* P < 0.001; **F,** b: Col-0 vs. PILS6 OX P < 0.05; PILS6 OX vs. *hrd1a hrd1b* P < 0.01; c: hrd1a hrd1b vs. PILS6 OX x *hrd1a hrd1b* P < 0.05). In all panels with boxplots: Box limits represent the 25th percentile and 75th percentile; the horizontal line represents the median. Whiskers display min. to max. values. All experiments were repeated at least three times.

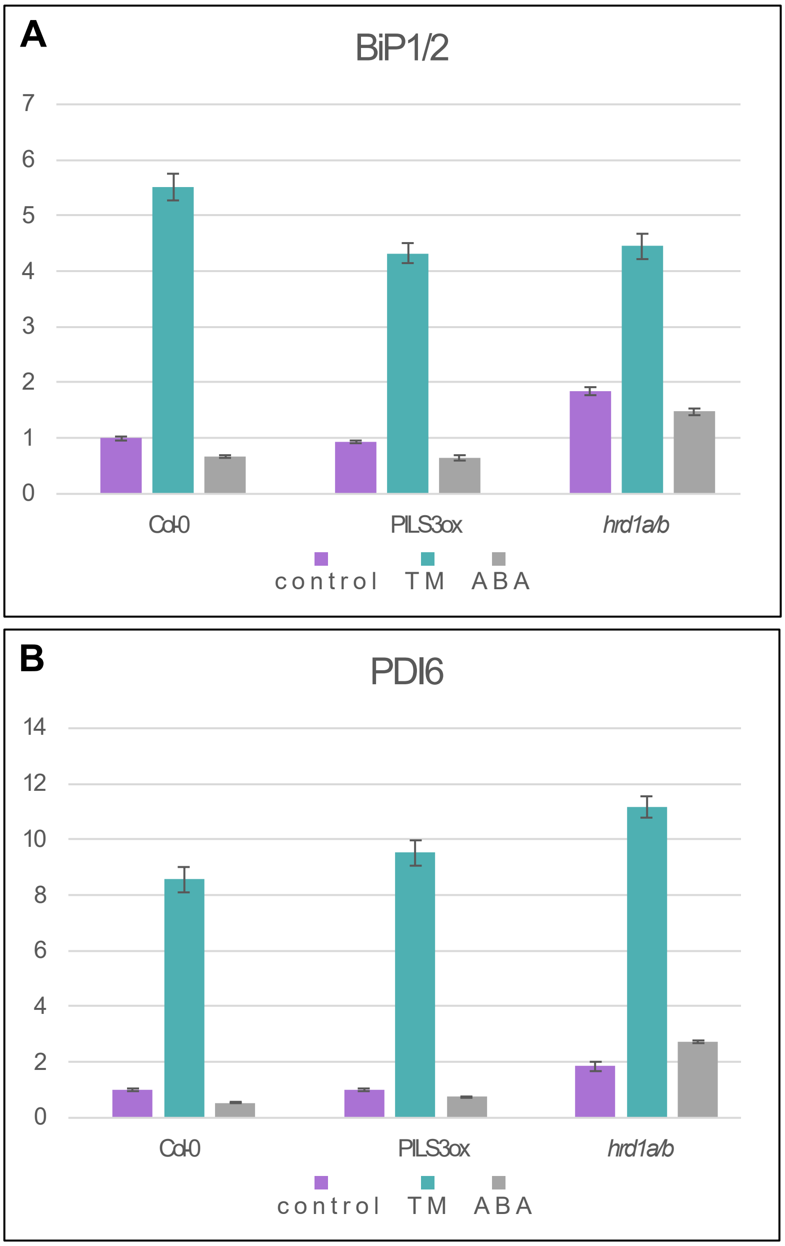


**Figure S5: ABA treatment does not induce ER stress.**
qPCR analysis detecting transcript levels of *BIP1, BIP2*, **(A)** and *PDI6* **(B)** normalized against *UBQ5* and *EIF4*. 4-day-old seedlings were treated for 4 hours with DMSO or 100 nM ABA or 5 µg/ml TM. Bars represent means ± SD, n = 3.

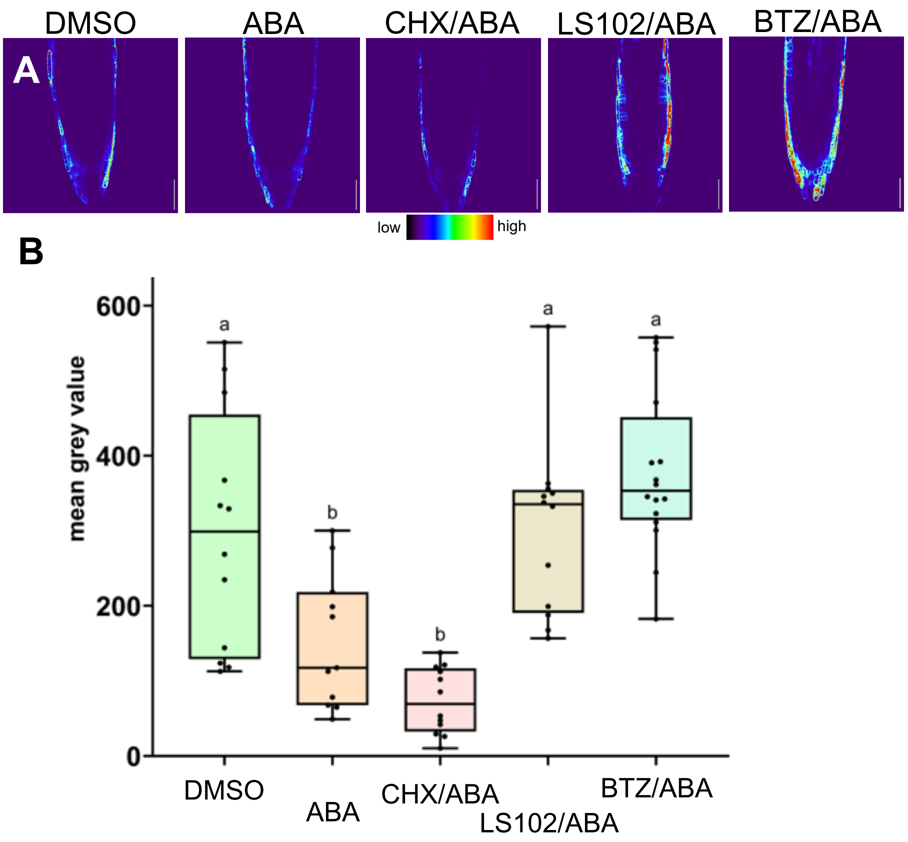


**Figure S6: Proteasome and HRD1 activity is required for the ABA-induced degradation of PILS3.**
**A-B,** Representative images **(A)** and quantifications **(B)** of GFP-PILS3, signal in 4-day seedlings. Seedlings were grown on solid ½ MS and treated with DMSO and 100 nM ABA as well as with ABA in combination with either 100 µM Cyclohecamide (CHX), 10 µM LS102, or 25 µM BTZ, in liquid ½ MS for 4h. Scale bars, 50 µm. n = 9-15 from seedling replicates pooled, one-way ANOVA followed by Tukey's multiple comparisons between treatments. In all panels with boxplots: Box limits represent 25th percentile and 75th percentile; horizontal line represents median. Whiskers display min. to max. values. P-Values: * $P < 0.05$, ** $P < 0.01$, *** $P < 0.001$, **** $P < 0.0001$. All experiments were repeated at least three times.
